# A highly Ca^2+^-permeable channelrhodopsin from the ancyromonad *Nutomonas longa* enables optogenetic control of Ca^2+^ signaling

**DOI:** 10.64898/2026.07.22.740102

**Authors:** Elena G. Govorunova, Yueyang Gou, Alex J. McDonald, Oleg A. Sineshchekov, Hai Li, Yumei Wang, François St-Pierre, Mingshan Xue, John L. Spudich

**Author notes:** These authors contributed equally to this work (Elena G. Govorunova, Yueyang Gou, Alex J. McDonald). Correspondence should be addressed to John L. Spudich.

## Abstract

Ca²⁺ is a ubiquitous regulator of cellular function, linking electrical activity to gene expression, secretion, metabolism, and synaptic plasticity. Yet, tools for its direct, time-resolved optical manipulation remain limited. Here, we report that *Nl*CCR, a channelrhodopsin from *Nutomonas longa,* possesses high Ca²⁺ permeability, enabling precise optical control of Ca²⁺ signaling. Compared with CapChR2, the most potent engineered Ca²⁺-conducting channelrhodopsin, *Nl*CCR combines larger and faster photocurrents, higher Ca²⁺ permeability, weaker desensitization, and reduced inward rectification. Mutational analysis identified determinants of Ca²⁺ selectivity and further enhanced it by introducing carboxylate residues at the channel’s central gate. *Nl*CCR’s blue-shifted absorption (445 nm) minimized optical crosstalk with a red-shifted Ca²⁺ indicator, laying the groundwork for all-optical experiments. In mouse cortical pyramidal neurons, *Nl*CCR enabled synaptic transmission independently of endogenous voltage-gated Ca²⁺ channels. These findings establish *Nl*CCR as a broadly applicable tool for direct, temporally precise manipulation of Ca²⁺-dependent signaling in living systems.

## Main text

Ca^2+^ is a ubiquitous intracellular messenger that regulates a vast array of cellular processes, including gene expression, neurotransmission, muscle contraction, secretion, fertilization, cell motility, and apoptosis. Genetically encoded fluorescent indicators are widely available for monitoring changes in intracellular Ca^2+^ concentration^1^. In contrast, genetically encoded tools for controlling Ca^2+^, based on fusing plant-derived light-sensitive domains to mammalian store-operated Ca^2+^ entry channels^2^ or their protein regulators^3^, suffer from limited responsiveness and low temporal resolution.

Channelrhodopsins (ChRs) are light-gated ion channels that guide phototaxis in protists, as demonstrated in the model chlorophyte alga *Chlamydomonas reinhardtii*^4^. When ChR genes are expressed in heterologous systems and activated by illumination, they generate transmembrane photocurrents that alter the membrane potential^5^. Under physiological conditions in mature mammalian neurons, photoactivation of cation ChRs (CCRs) leads to H^+^ and Na^+^ influx^5^; anion ChRs (ACRs), to Cl^-^ influx^6^; and kalium ChRs (KCRs), to K^+^ efflux^7^. The technique known as optogenetics^8^ uses CCRs to photoactivate neurons and ACRs and KCRs to photoinhibit them. The late component of photoreceptor current in *C. reinhardtii*, which predominates at low stimulus intensities, is primarily carried by Ca^2+9^. However, the permeability of heterologously expressed wild-type *C. reinhardtii* ChR2 (*Cr*ChR2) to Ca^2+^ relative to Na^+^ (P_Ca_/P_Na_) is only 0.15^10^. *Cr*ChR1, the other CCR from the same organism, is even less permeable to Ca^2+11^. We hypothesized that the late photoreceptor current in *C. reinhardtii* is mediated by secondary Ca^2+^ channels activated upon ChR photoexcitation^9, 12^, but their genes had not yet been cloned.

Studies on heterologously expressed *Cr*ChR2, the most popular excitatory optogenetic tool, identified several mutations increasing Ca^2+^ permeation, culminating in the creation of CapChR1 (Ca^2+^-permeable ChR1; *Cr*ChR2_S63D_L132C_T159C_N258E)^13^. A corresponding quadruple mutant of *Chloromonas oogama* ChR (*Co*ChR), named CapChR2, exhibited even larger relative permeability to Ca^2+^ than CapChR1. Both CapChRs mediated photoinduced Ca^2+^ influx in mammalian neurons and *Drosophila* brain explants^13^. However, CapChR2 exhibits slow photocurrent decay and strong inward rectification, which limits its temporal precision and restricts Ca^2+^ influx, respectively. Moreover, CapChR2 retains substantial sensitivity to yellow light, hindering its spectral compatibility with many red-emitting indicators in all-optical experiments. Together, these limitations constrain the use of CapChR2 across a broad range of applications requiring precise manipulation of intracellular Ca²⁺ dynamics.

These limitations prompted us to ask whether superior Ca²⁺-conducting channelrhodopsins might already exist in nature. As we predicted, we found such a channel in a ChR family from ancient bacterivorous ancyromonad flagellates ^14^. Two representatives of this family, *Ancyromonas sigmoides* (*Ans*) ACR and *Fabononas tropica* (*Ft*) ACR, are anion-selective. In contrast, the homolog from *Nutomonas longa* (*Nl*CCR) does not conduct anions. Here, we show that *Nl*CCR is highly permeable to Ca^2+^ and outperforms CapChR2 across several other biophysical parameters relevant to optogenetic applications. Using a spectrally compatible Ca^2+^ fluorescent indicator, we confirm that photoactivation of *Nl*CCR results in a greater increase in intracellular Ca²⁺ concentration than that of CapChR2 and displays negligible optical crosstalk. Finally, we show that *Nl*CCR expression in cortical pyramidal neurons enables light-driven synaptic transmission, allowing optical modulation of Ca²⁺ without eliciting action potentials.

## Results

### *Nl*CCR is a robust Ca^2+^-permeable channel

We sought to characterize *Nl*CCR and compare it with CapChR2, the most potent of the previously available Ca^2+^-permeable ChR variants^13^. We fused these ChRs to mCherry at their C-terminus for visualization, expressed them in human embryonic kidney (HEK293) cells, and recorded photocurrents using the high-throughput, planar, automated patch-clamp platform SyncroPatch 384 ^15^. We used 4-hole chips to increase the probability of capturing transgene-expressing cells in each well. We also included *Cr*ChR2 in this experiment to test whether our assay could detect the previously reported difference in selectivity between CapChR2 and *Cr*ChR2^13^. Figure 1a shows representative photocurrent traces recorded upon incremental holding voltages using the physiological, Na^+^-based external solution containing 2 mM Ca^2+^ (black), and after the replacement of all external Na^+^ with Ca^2+^ and repeating the measurements in the same cells (red; see Methods and Supplementary Table 1 for the solution compositions and other details). Photocurrent–voltage relationships in both solutions revealed strong, modest, and minimal inward rectification in CapChR2, *Cr*ChR2, and *Nl*CCR, respectively (Figure 1b).

**Figure 1.**
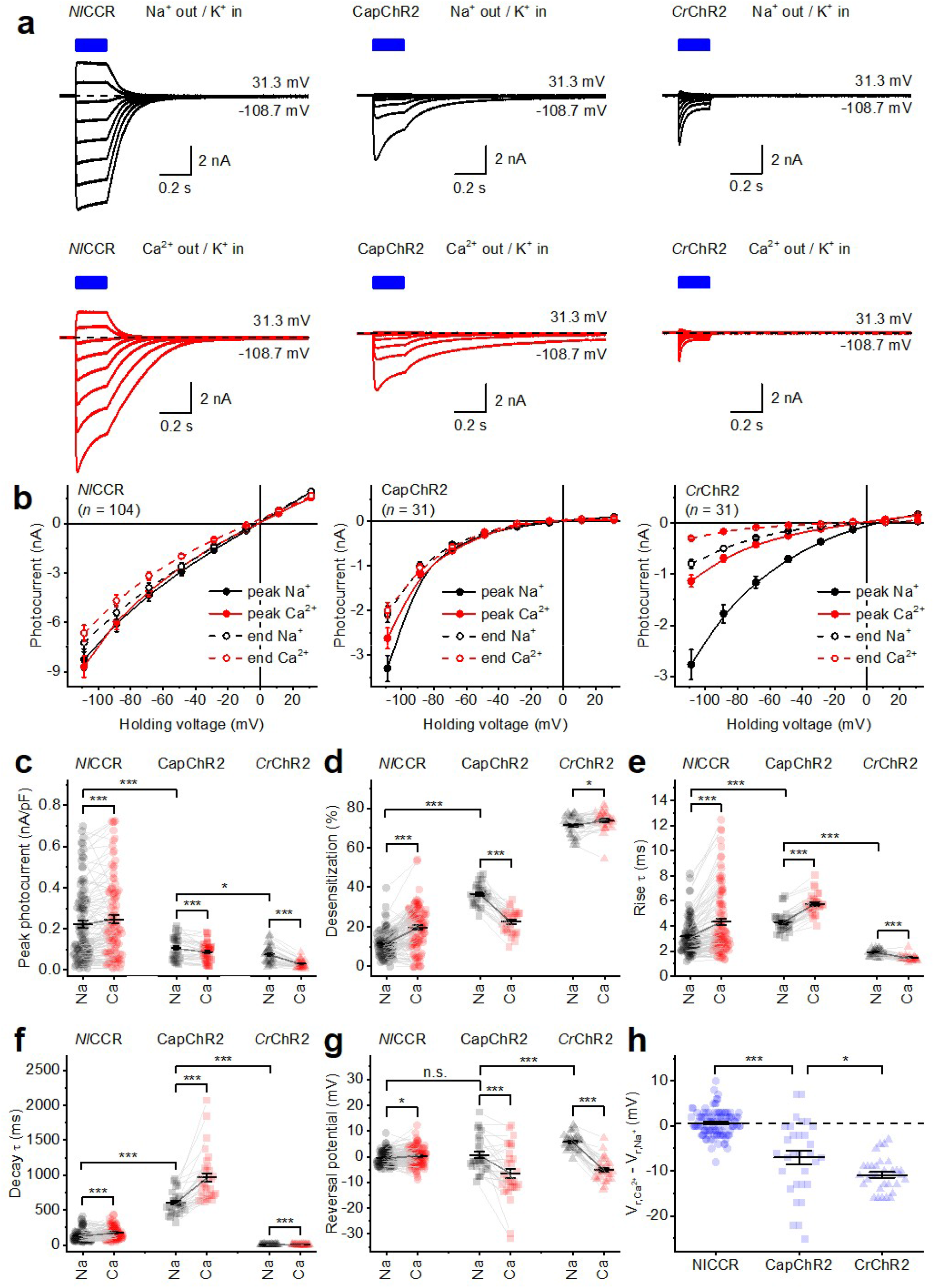
Comparative analysis of *Nl*CCR, CapChR2, and *Cr*ChR2 by automated patch clamping. **a**, Representative photocurrent traces at indicated holding voltages in response to 200-ms light pulses (blue bars), recorded using Na^+^- (black) or Ca^2+^-based (red) external solutions. Holding voltages were corrected for LJP. **b**, The voltage dependencies of the photocurrent measured at peak (filled symbols) and the end of illumination (empty symbols) in the Na^+^-based external solution (black) and after its exchange to the Ca^2+^-based external solution (red). The symbols represent mean ± SEM values; the numbers of wells sampled per variant (n) are shown in the plots. **c**, Peak photocurrent amplitudes at -108.7 mV, normalized to the membrane capacitance (a proxy for the membrane area). **d**, Desensitization during 200-ms light pulses, quantified as the decrease in photocurrent from its peak to the end of illumination. **e,f**, Photocurrent rise (e) and decay (f) time constants (τ) at -108.7 mV. **g**, V_r_ values calculated at the peak photocurrent time. **h**, V_r_ shifts after the replacement of extracellular Na^+^ with Ca^2+^. In b-g, the black symbols show the data obtained in the Na^+^-based external solution, and the red symbols show the data obtained in the Ca^2+^-based external solution. In c-h, the symbols represent the individual well data; the lines and error bars represent the mean ± SEM. *, P < 0.05; **, P < 0.01; ***, P < 0.005 by the two-tailed Wilcoxon signed-rank test for comparisons between different solutions for the same ChR and the two-tailed Mann-Whitney test for comparisons between different ChRs. The raw numerical data, the numbers of wells sampled (n), and the exact p-values are provided in the Source Data File.

At -108.7 mV (adjusted for the liquid junction potential, LJP), the mean *Nl*CCR photocurrent response per pF to the first light pulse was more than twice larger than that of CapChR2 (Fig. 1c). This advantage was observed despite the use of 470-nm illumination, which is close to the absorption maximum (λ_max_) of CapChR2^13^ but activates *Nl*CCR (λ_max_ 445 nm) at only ∼75% of its maximal efficiency ^14^. *Nl*CCR’s advantage was even larger at more depolarized voltages (∼6 times at -70 mV) due to CapChR2’s strong inward rectification (Fig. 1b). In the physiological external solution, *Nl*CCR showed less desensitization (photocurrent reduction during illumination) than CapChR2 (Fig. 1d, black). Interestingly, high extracellular Ca^2+^ increased desensitization of *Nl*CCR and *Cr*ChR2, but decreased it in CapChR (Fig. 1d, red), suggesting that desensitization in the two ChRs is brought about by different photochemical processes.

Fast ChR kinetics are essential for applications requiring high temporal precision, including the optical reproduction of rapid neuronal signals. Both photocurrent rise (τ 3.1 vs. 4.3 ms in the physiological solution) and decay (135 vs. 606 ms) of *Nl*CCR, estimated by monoexponential approximation, were faster than those of CapChR2 (Figs. 1e,f). The reversal potentials (V_r_) of *Nl*CCR and CapChR2 in the Na^+^ solution were close to zero, indicating K^+^/Na^+^ permeability ratios close to one (Fig. 1g). Replacing extracellular Na^+^ with Ca^2+^ caused a small but statistically significant (see Source Data File for statistical analysis) positive V_r_ shift for *Nl*CCR and to negative V_r_ shifts for the other two ChRs (Fig. 1g,h), reflecting the higher relative Ca^2+^ permeability of *Nl*CCR. *Cr*ChR2 showed a larger negative V_r_ shift than CapChR2, consistent with its lower relative Ca^2+^ permeability, as previously observed using manual patch-clamping ^13^. Taken together, these findings demonstrate *Nl*CCR’s superior biophysical characteristics compared with those of CapChR2.

### *Nl*CCR mediates photoinduced Ca^2+^ influx under physiological conditions

We next sought to quantify *Nl*CCR-mediated Ca²⁺ influx in the presence of physiological extracellular ions, including Na⁺, and to evaluate its compatibility with all-optical Ca²⁺ manipulation and imaging. We selected the red genetically encoded Ca²⁺ indicator PinkyCaMP^16^, which is spectrally compatible with blue-light activation of *Nl*CCR (Fig. 2a). The assay employed brief 10-ms pulses of 440/20-nm light (peak wavelength/bandwidth) to activate ChRs and continuous 575/25-nm illumination to monitor PinkyCaMP fluorescence (Fig. 2b).

**Figure 2.**
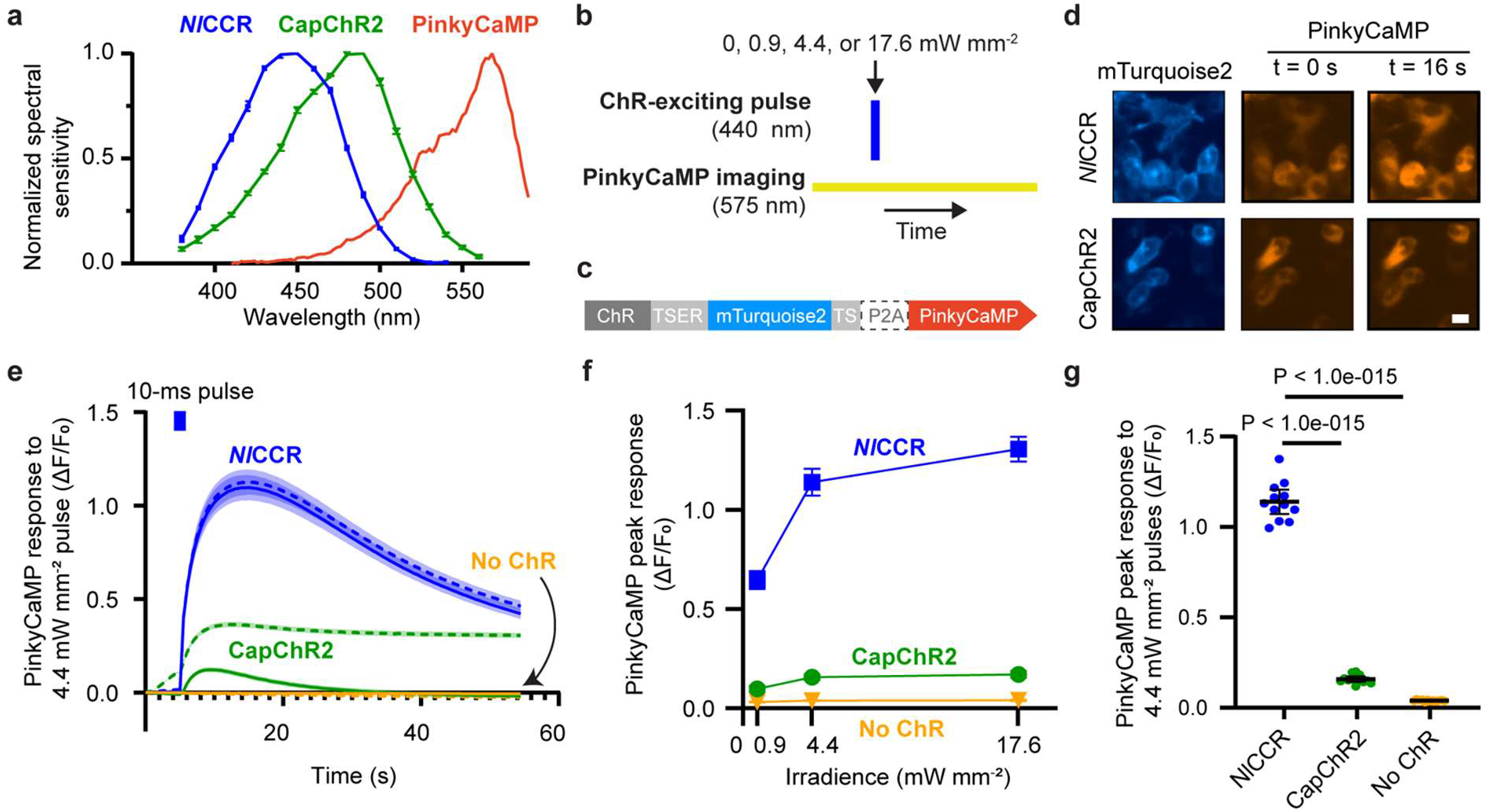
Fluorescence imaging of Ca^2+^ influx generated by photoactivation of *Nl*CCR and CapChR2 a, Action spectra of *Nl*CCR and CapChR2 photocurrents, and fluorescence excitation spectrum of PinkyCaMP. All spectra are normalized to peak values. Data points: mean values. Error bars: SEM. n = 9 (*Nl*CCR) and 10 (CapChR2) cells. **b,** All-optical illumination protocol. PinkyCaMP fluorescence was recorded with 575/25-nm light for 55 s while a single 10-ms pulse of 440/20-nm light was delivered at t = 5 s. Pulses of different irradiances were delivered to separate fields of view. **c,** Schematic representation of the bicistronic plasmid-based cassette co-expressing ChR and PinkyCaMP using a ribosome-skipping peptide (P2A). TSER: trafficking sequences promoting export from the Golgi (TS) and the endoplasmic reticulum (ER). ChRs were fused to the CFP mTurquoise2. **d,** Representative groups of HEK293-Kir2.1 cells. *Left*: ChR-mTurquoise2 (mT2) fluorescence. *Right*: PinkyCaMP fluorescence at t = 0 or 16 s, near the time of peak response. **e-g,** Fluorescence imaging of Ca^2+^ transients following ChR photoactivation. Responses in the absence of ChRs (“No ChR”) are also shown. n = 12 independently transfected wells. **e.** PinkyCaMP responses (dashed lines). Traces were corrected for optical crosstalk by subtracting the corresponding recordings collected without the 440/20-nm pulse (solid lines). Due to spectral bleed-through of mTurquoise2, the frame during which the 440/20-nm light was pulsed is omitted from the traces. Shaded areas: 95% confidence interval (CI). **f,** Dependence of PinkyCaMP peak responses on blue-light irradiance. Error bars: 95% CI (n = 12 wells for all conditions except *Nl*CCR at 17.6 mW mm^-2^, for which n = 11). **g.** Statistical comparisons of PinkyCaMP responses to 4.4-mW mm^-2^ pulses of light. P-values: Dunnett’s test.

To enable simultaneous expression of both proteins, we generated plasmids co-expressing ChRs and PinkyCaMP (Fig. 2c). ChRs were fused to Golgi-export (TS)^17^ and endoplasmic reticulum-export (ER)^18, 19^ trafficking motifs to enhance plasma membrane localization and to the cyan fluorescent protein mTurquoise2^20^ to monitor expression and subcellular distribution. A control construct expressing PinkyCaMP alone was generated in parallel.

All plasmids were transiently expressed in HEK293-Kir2.1 cells, which maintain a physiologically relevant resting membrane potential (∼−70 mV) and therefore permit quantitative comparisons under neuron-like conditions^21, 22^. Both *Nl*CCR and CapChR2 showed prominent plasma membrane fluorescence, as visualized by mTurquoise2, along with intracellular fluorescence likely corresponding to proteins localized to the endoplasmic reticulum and Golgi apparatus, a common feature of ChR expression (Fig. 2d, blue). As expected, PinkyCaMP was localized to the cytoplasm and reported increases in intracellular Ca²⁺ as increased red fluorescence (Fig. 2d, orange).

CapChR2 produced a gradual increase in PinkyCaMP fluorescence before blue-light stimulation (Fig. 2e, dashed lines), which we attribute to partial activation by the 575/25 nm imaging illumination. Although CapChR2 displays relatively low sensitivity to wavelengths above 550 nm, this residual sensitivity is sufficient to induce measurable activation under standard imaging conditions (Fig. 2a). In contrast, under the same imaging conditions, *Nl*CCR elicited detectable Ca²⁺ transients only in response to blue-light stimulation (Fig. 2e, blue dashed line), consistent with its more blue-shifted action spectrum (Fig. 2a). To enable direct comparison of the two ChRs, fluorescence traces were corrected for optical crosstalk by subtracting responses recorded during 575/25-nm imaging in the absence of 440/20-nm stimulation (Fig. 2e, solid lines).

*Nl*CCR generated markedly larger Ca²⁺ transients than CapChR2 across all tested irradiances (Figs. 2f,g). Peak responses increased with light intensity and approached saturation at higher irradiances, consistent with limits set by channel activation (Fig. 2f; Extended Data Fig. 1). At 4.4 mW mm⁻², *Nl*CCR elicited a ∼7-fold larger Ca²⁺ response than CapChR2 (Fig. 2g), consistent with its ∼6-fold larger photocurrent at -70 mV (Fig. 1b). Together, these results demonstrate that *Nl*CCR enables efficient, stimulus-locked Ca²⁺ influx and spectral compatibility with the red indicator PinkyCaMP. These properties support all-optical manipulation and monitoring of intracellular Ca²⁺ dynamics with reduced spectral interference and substantially greater efficacy than previously engineered Ca²⁺-conducting channelrhodopsins.

### Activation of *Nl*CCR induces synaptic transmission independently of endogenous voltage- gated Ca^2+^ channels

To characterize and validate the action of *Nl*CCR in neurons, we evaluated whether activation of *Nl*CCR could conduct enough Ca²⁺ to directly trigger synaptic transmission, bypassing action potential generation and endogenous voltage-gated Ca^2+^ channel (VGCC) activation. We selectively introduced a plasmid expressing *Nl*CCR fused to EYFP and the membrane-targeting motif Kv2.1C-linker-TlcnC^23^ (*Nl*CCR-EYFP-Kv2.1C-linker-TlcnC, referred to as *Nl*CCR from now on) and another plasmid expressing the red fluorescent protein tdTomato into layer 2/3 pyramidal neurons of the mouse somatosensory cortex by *in utero* electroporation. Acute brain slices prepared from 3–6-week-old mice exhibited EYFP fluorescence in layer 2/3, where the transfected cell bodies resided, and layer 5, to which layer 2/3 neurons projected their axons (Fig. 3a), confirming *Nl*CCR expression in the somatodendritic domains and the axons, respectively. For comparison, CapChR2-EYFP-Kv2.1C-linker-TlcnC (CapChR2) and *Cr*ChR2_H134R-EYFP (*Cr*ChR2_H134R), a widely used CCR variant with minimal Ca^2+^ permeability^24^, were expressed using the same approach.

**Figure 3.**
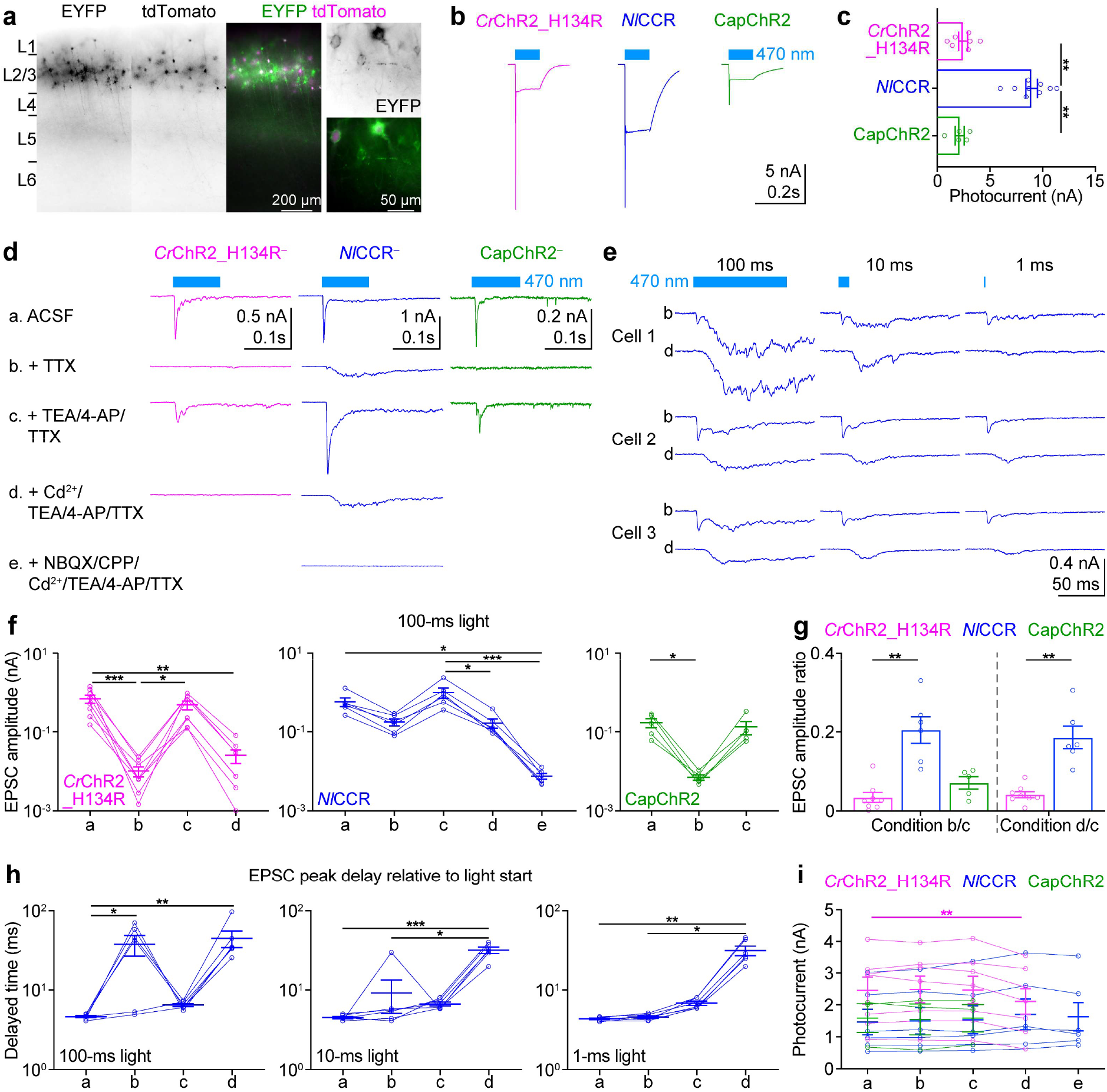
*Nl*CCR generates larger photocurrents than CapChR2 and *Cr*ChR2_H134R and enables neurotransmitter release without activating VGCCs. (**a**) Representative overview (left) and high-magnification (right-most) fluorescence images of 300-µm-thick brain slices expressing tdTomato and *Nl*CCR-EYFP-Kv2.1C-linker-TlcnC (*Nl*CCR) in cortical layer 2/3 pyramidal neurons. The axons of *Nl*CCR^+^ pyramidal neurons in layer 2/3 ramify in layer 5. L, layer. (**b,c**) Representative photocurrent traces (b) and summary data (c) of photocurrent amplitudes at the end of a 100-ms 470-nm light pulse (38.7 mW mm^-2^) from *Cr*ChR2_H134R- EYFP (*Cr*ChR2_H134R, *n* = 7)-, *Nl*CCR (*n* = 9)-, or CapChR2-EYFP-Kv2.1C-linker-TlcnC (CapChR2, *n* = 5)-expressing neurons. (**d**) Representative current traces of *Cr*ChR2_H134R^−^, *Nl*CCR^−^, and CapChR2^−^ neurons recorded at -60 mV under indicated conditions a, b, c, d, or e, in response to 100-ms pulses of 470-nm light (38.7 mW mm^-2^). (**e**) Example current traces from three *Nl*CCR^−^ neurons recorded at -60 mV under the condition b or d, in response to a 100-ms, 10-ms or 1-ms 470-nm light pulse (38.7 mW mm^-2^). (**f**) Summary data of evoked EPSC peak amplitudes of recorded ChR^−^ neurons in the brain slices expressing *Cr*ChR2_H134R (*n* = 8), *Nl*CCR (*n* = 6), or CapChR2 (*n* = 5) in response to 100-ms 470-nm light stimulation under the indicated conditions in d. (**g**) Ratios of evoked EPSC peak amplitudes in TTX vs. TEA/4-AP/TTX (left) and in Cd^2+^/TEA/4-AP/TTX vs. TEA/4-AP/TTX (right) from **f**. (**h**) Summary data of EPSC peak delays after the light onset of recorded ChR^−^ neurons in the brain slices expressing *Nl*CCR (*n* = 6) in response to 100-ms, 10-ms, and 1-ms illumination. (**i**) Photocurrent amplitude at the end of a 100-ms 470-nm light pulse from *Cr*ChR2_H134R (*n* = 7)-, *Nl*CCR (*n* = 6)-, or CapChR2 (*n* = 3)-expressing neurons under the indicated conditions in d (38.7 mW mm^-2^ for *Cr*ChR2_H134R and CapChR2 or 1.8 µW mm^-2^ for *Nl*CCR). In all panels, each symbol represents a recorded neuron. Magenta, blue, and green indicate data from the mice expressing *Cr*ChR2_H134R (two males), *Nl*CCR (two females), and CapChR2 (one female), respectively. Summary data are expressed as mean ± SEM. *, P < 0.05; **, P, < 0.01; ***, P < 0.001 by Friedman test with Dunn’s correction (c, f, g left, h and i) or two-tailed Mann-Whitney test (g right).

To quantify photocurrents, we performed whole-cell voltage-clamp recordings from ChR-expressing (ChR⁺) neurons in acute brain slices using a K⁺-based internal solution. Photostimulation with 100-ms pulses of 470-nm light could elicit unclamped action potentials at light onset in neurons expressing each of the three ChRs (Fig. 3b). However, *Nl*CCR generated much larger photocurrent than CapChR2 and *Cr*ChR2_H134R (Fig. 3c), which may be due to its larger channel conductance, better expression on the plasma membrane, or both.

To evaluate whether these tools can support light-induced neurotransmission, we stimulated ChRs with 100-ms pulses of 470-nm light and recorded excitatory postsynaptic currents (EPSCs) from neighboring ChR-negative (ChR^−^) layer 2/3 pyramidal neurons using whole-cell voltage-clamp with a Cs⁺-based internal solution. As expected, activation of *Cr*ChR2_H134R elicited fast EPSCs (Fig. 3d, condition a) that depended on presynaptic action potentials, as demonstrated by their abolition by the voltage-gated Na^+^ channel blocker tetrodotoxin (TTX) (Fig. 3d, condition b). Blocking voltage-gated K^+^ channels using tetraethylammonium (TEA) and 4-aminopyridine (4-AP) restored evoked EPSCs (Fig. 3d, condition c), consistent with enhanced presynaptic depolarization ^25^ and subsequent activation of voltage-gated Ca²⁺ channels (VGCCs). Accordingly, EPSCs were again eliminated by the VGCC blocker Cd²^+^ (Fig. 3d, condition d).

Next, we performed similar experiments with *Nl*CCR and CapChR2. Activation of *Nl*CCR resulted in not only fast EPSCs but also slower inward currents (Fig. 3d, condition a). TTX drastically reduced, but did not completely eliminate, fast EPSCs, as small fast EPSCs were still often observed at the onset of light stimulation (Fig. 3d,e, condition b). Importantly, the slower inward currents persisted (Fig. 3d,e, condition b). The fast EPSCs were restored by TEA/4-AP (Fig. 3d, condition c) and abolished by Cd^2+^ (Fig. 3d,e, condition d), as for *Cr*ChR2_H134R. However, the slower inward currents were not affected by Cd^2+^ (Fig. 3d,e, condition d). We confirmed that these inward currents were EPSCs, as they were abolished by the glutamate receptor antagonists NBQX and CPP (Fig. 3d, condition e). The residual fast EPSCs evoked by *Nl*CCR in the presence of TTX were sensitive to Cd^2+^, indicating that even without TEA/4-AP, the large *Nl*CCR photocurrents can modestly open VGCCs to trigger neurotransmitter release. In the absence of Ca^2+^ influx through the VGCCs, *Cr*ChR2_H134R could not evoke neurotransmission, whereas *Nl*CCR mediated sufficient Ca^2+^ influx to trigger neurotransmitter release (Fig. 3f). In contrast, activation of CapChR2 induced very little neurotransmitter release in the presence of TTX (Fig. 3d, condition b; Fig. 3f), although TEA/4-AP also recovered the fast EPSCs (Fig. 3d, condition c; Fig. 3f). We could not test the effect of blocking VGCCs because Cd^2+^ unexpectedly activated CapChR2 in the absence of light stimulation.

The amplitudes of both fast EPSCs and slow EPSCs depend on the number of ChR^+^ neurons and photocurrents in ChR^+^ neurons, both of which vary across preparations. To control for these variabilities, we compared the of EPSC peak ratios in the Cd^2+^/TEA/4-AP/TTX and TEA/4-AP/TTX conditions for *Cr*ChR2_H134R and *Nl*CCR, as these ratios reflect the fraction of Ca^2+^ influx through ChRs. Similarly, we also compared the EPSC peak ratios in the TTX and TEA/4-AP/TTX conditions for *Cr*ChR2_H134R, *Nl*CCR, and CapChR2. Both ratios were significantly larger higher for *Nl*CCR than for *Cr*ChR2_H134R and CapChR2 (Fig. 3g), indicating that *Nl*CCR activation mediates much more Ca^2+^ influx in neurons than *Cr*ChR2_H134R and CapChR2.

We next examined the effects of shorter photostimulation durations (10 ms and 1 ms) on ChR-mediated synaptic transmission. Whereas fast EPSC amplitudes were similar across all three stimulation durations, slow EPSCs scaled strongly with pulse duration (Fig. 3f and Extended Data Fig. 2a). In the presence of TTX, 10-ms and 1-ms stimuli produced slow EPSCs whose magnitude often remained below that of the residual fast EPSCs, resulting in early peaks (Fig. 3h). By contrast, 100-ms stimulation produced larger slow EPSCs that exceeded the residual fast component in most neurons, shifting the EPSC peak to later times (Fig. 3e,h, condition b). In the presence of Cd²⁺/TEA/4-AP/TTX, EPSC peaks were delayed for all stimulation durations because the residual fast EPSC component was eliminated, leaving only the slow responses (Fig. 3h).

Finally, we confirmed that all pharmacological treatments caused negligible changes in the photocurrents of the tested ChRs, ruling out altered ChR activity as an explanation for the observed differences in synaptic transmission. (Fig. 3l). Together, these results show that activation of *Nl*CCR in neurons generates sufficient Ca^2+^ influx at the axonal terminals to directly induce synaptic transmission independently of action potentials and endogenous VGCCs.

### Amino acid residues responsible for *Nl*CCR Ca^2+^ permeability

Having established the utility of *Nl*CCR for manipulating intracellular Ca²⁺ in intact brain tissue, we next sought to define the molecular determinants of its high Ca²⁺ permeability and evaluate its potential as a scaffold for engineering next-generation variants with further enhanced Ca²⁺ permeability and other useful properties. Among all known ChRs, the *Nl*CCR protein sequence is most closely related to other ancyromonad ChRs (34/52% and 31/49% identity/similarity with *Ans*ACR and *Ft*ACR, respectively), both of which transport anions^14^. However, the *Nl*CCR sequence differs from the ancyromonad ACRs at residue positions critical for the transport activity in microbial rhodopsins (Extended Data Fig. 3). At the primary proton acceptor position, corresponding to Asp85 in *Halobacterium salinarum* bacteriorhodopsin (*Hs*BR), *Nl*CCR exhibits Phe instead of Gly found in both ancyromonad ACRs. The residue corresponding to *Hs*BR’s Thr89 is replaced with Ser in *Nl*CCR, but is conserved in both ancyromonad ACRs. The proton-donor residue (Asp96 in *Hs*BR) is conserved in *Nl*CCR but replaced by Ala in the ancyromonad ACRs. At the position of *Hs*BR’s Leu99, which shield Asp96 from the aqueous cytoplasmic environment, *Nl*CCR exhibits Lys, in contrast to Gln in the ancyromonad ACRs. The second photoactive-site carboxylate (Asp212 in *Hs*BR) is replaced with Glu in *Nl*CCR but conserved in the ancyromonad ACRs. Extended Data Figure 4a shows the positions of these residues in the homology models, as no molecular structures of ancyromonad ChRs and CapChR2 have been obtained to date. We individually replaced these *Nl*CCR residues with the corresponding residues in the ancyromonad ACRs (the F85G, S89T, D96A, K99N, and E233D mutations) to test their potential roles in Ca^2+^ permeability. CapChR2 was engineered from the wild-type *Co*ChR by combining four point mutations^13^. The L112C and T139C mutations in TM3 and TM4, respectively, were made because the corresponding mutations in *Cr*ChR2 (L132C and T159C) promoted retinal binding and plasma membrane targeting, and reduced desensitization, respectively ^26^. In all three ancyromonad ChRs, Leu112/132 is replaced with Val (Val94 in *Nl*CCR) (Extended Data Fig. 3). TM4 of ancyromonad ChRs is very divergent from that of chlorophyte CCRs (Extended Data Fig. 3), but the structural alignment of the homology models suggests that Met141 in *Nl*CCR corresponds to Thr139 in CapChR2 (Extended Data Fig. 4b). The other two mutations made to create CapChR2 (S43D in TM1 and N238E in TM7) introduced carboxylate residues into the so-called “central gate”, a network of H-bonded residues centered on Glu70 (corresponding to Glu90 in *Cr*ChR2) in TM2 near the Schiff base. The original Ser and Gln residues are conserved in *Nl*CCR (Ser21 and Gln238, respectively), although Glu70 is replaced with Ala48 (Extended Data Fig. 4b). CapChR2 exhibits a Glu residue (Glu103) in the position of *Hs*BR’s Asp85, as is typical of most chlorophyte CCRs. It differs from wild-type *Cr*ChR2 by two carboxylated residues (Asp83 and Asp84) in the TM2-TM3 loop, the second of which is replaced with Asn62 in *Nl*CCR (Extended Data Fig. 3). We mutated *Nl*CCR residues to the residues found at the corresponding positions in CapChR2 (S21D, A48E, N62D, F85E, V94C, M141C, and N238E).

We tested both sets of mutants using automated patch-clamp recordings, as described above. Extended Data Figure 5 shows representative photocurrent traces recorded from the mutants at incremental voltages, and Extended Data Figure 6 shows the current-voltage dependencies. In the physiological, Na^+^-based external solution, the V_r_ values in all tested mutants except A48E were indistinguishable from the wild-type value by one-way ANOVA followed by the Tukey test (Fig. 4a, black; see Source Data for exact p values), suggesting the same Na^+^/K^+^ permeability ratio, whereas a slightly more positive V_r_ of A48E suggests a slightly higher permeability for Na^+^ than for K^+^.

**Figure 4.**
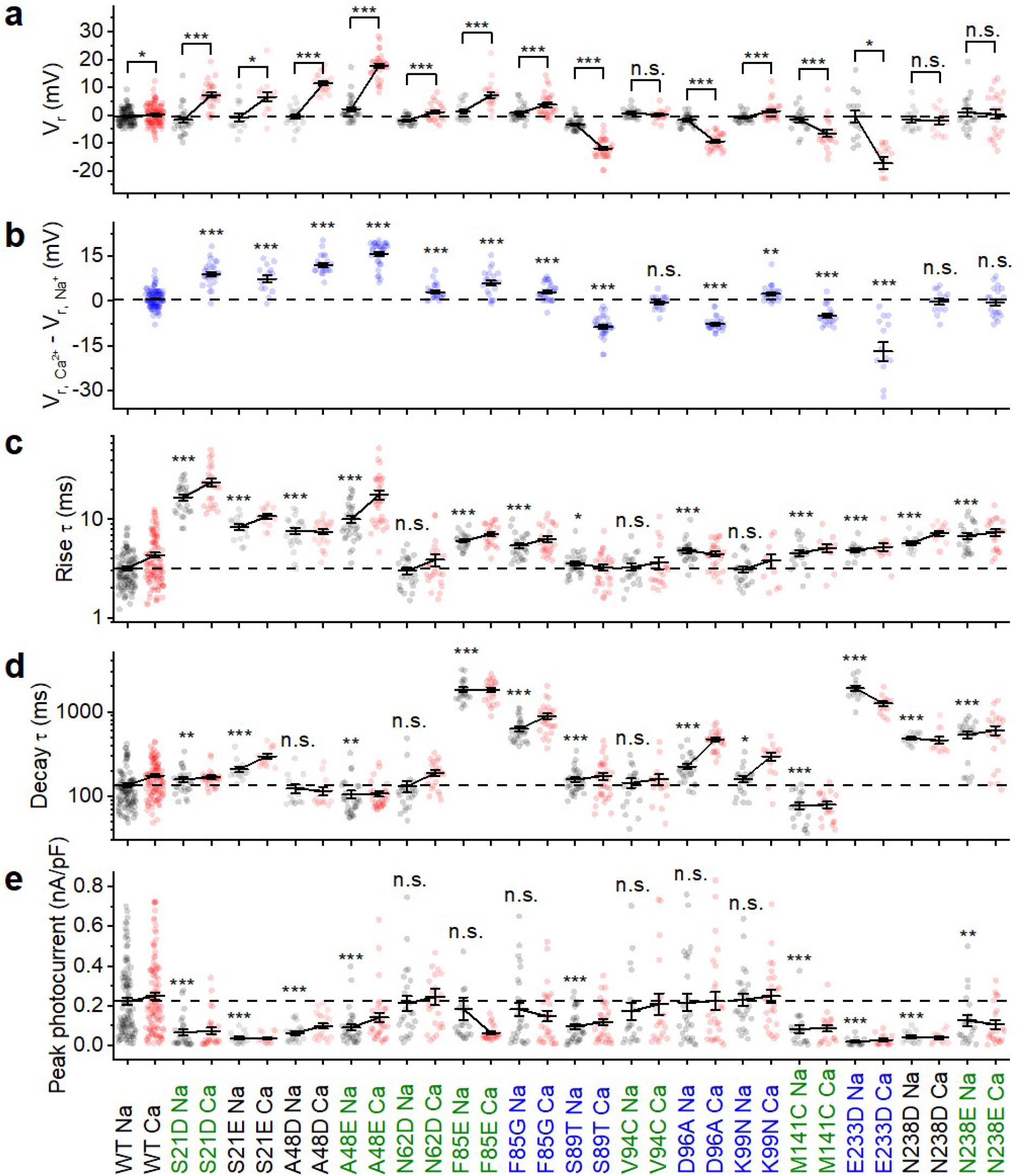
Mutant analysis of *Nl*CCR. **a**, Reversal potentials (V_r_) determined at the peak photocurrent time. **b**, V_r_ shifts after the replacement of extracellular Na^+^ with Ca^2+^. **c,d**, Time constants (τ) of the photocurrent rise (c) and decay (d) at -108.7 mV. **e**, Peak photocurrent amplitudes at -108.7 mV, normalized to the membrane capacitance. In a and c-e, the black symbols show the data obtained in the Na^+^-based external solution, and the red symbols show the data obtained in the Ca^2+^-based external solution. The symbols represent the individual well data; the solid lines represent the mean ± SEM values. *, P < 0.05; **, P < 0.01; ***, P < 0.005; n.s., non-significant by the two-tailed Wilcoxon signed-rank test for comparisons between the Na^+^ and Ca^2+^ conditions for the same variant (a), and by the two-tailed Mann-Whitney test for comparisons between the mutants and wild type in the Na^+^ conditions (b-e). The numerical values and the numbers of wells sampled (n) are provided in the Source Data. The dashed lines show the values obtained for the wild-type *Nl*CCR in the standard (Na^+^-based) solution (a and c-e), or the V_r_ shift in the wild-type *Nl*CCR (b).

Three of the five mutants, in which *Nl*CCR residues were replaced with *Ans*ACR/*Ft*ACR residues (S89T, D96A, and especially E233D), showed large negative V_r_ shifts upon replacement of extracellular Na^+^ with Ca^2+^ compared to the wild-type *Nl*CCR (Fig. 4a, red and Fig. 4b), indicating a strong reduction of Ca^2+^/Na^+^ relative permeability. Interestingly, in S89T and E233D, these negative shifts were combined with an increase in photocurrent at negative voltages (Extended Data Figure 6), revealing separate regulation of channel gating and selectivity. The reciprocal A97D mutation in *Ans*ACR, which introduces the aspartate found at the corresponding position in *Nl*CCR, did not confer Ca²⁺ permeability: the V_r_ shift measured upon replacement of Na^+^ with Ca^2+^ in the mutant did not change from that in the wild-type *Ans*ACR (-2.7 ± 0.6 and -2.5 ± 0.6 mV, mean ± SEM, n = 20 and 19 wells, respectively; p = 0.8 by the two-tailed Mann-Whitney test). However, *Ans*ACR_A97D showed a much slower photocurrent decay than the wild-type *Ans*ACR (Extended Data Fig. 7a,b), confirming the importance of this position for channel gating in ancyromonad ChRs. Remarkably, *Ans*ACR_A97D also conferred strong outward rectification (Extended Data Fig. 7c,d), consistent with an earlier observation that *Gt*ACR1 mutations that introduce a negative charge in the cytoplasmic part of the conduction pathway promote this phenotype, despite the low sequence homology between the two proteins^27^.

The *Nl*CCR-to-CapChR2 replacements, which introduced carboxylate side chains into the central gate (S21D and especially A48E), yielded positive V_r_ shifts, indicating increased Ca^2+^ permeability (Fig. 4a). Similar shifts were observed in the *Nl*CCR_S21E and A48D mutants, suggesting the importance of the side chain’s negative charge. However, neither aspartate nor glutamate replacement of the third central gate residue, Asn238, shifted the V_r_ to more positive values, in contrast to the N238E mutation of CapChR2^13^. Intriguingly, *Nl*CCR_S21D/E and A48D/E mutations, in addition to increasing Ca^2+^ permeability, showed much slower photocurrent rise than the wild-type *Nl*CCR (Fig. 4c). Replacement of Phe85 (corresponding to *Hs*BR’s Asp85) with either Glu, found in this position in CapChR, or Gly, found in *Ans/Ft*ACRs, strongly slowed the photocurrent decay, as did the E233D mutation of the second carboxylate in the photoactive site (Fig. 4d). Mutagenetic neutralization of the conserved Asp96, the proton donor to the Schiff base in *Hs*BR, also showed a pronounced effect on the photocurrent decay (Fig. 4d). The strongest suppression of the photocurrent amplitude was found in the S21E, E233D and N238D mutants, indicating the involvement of these residue positions in the channel’s conductance (Fig. 4e). Together, these findings identify residues that shape NlCCR Ca²⁺ permeability and demonstrate that this property can be further enhanced by engineering the central gate.

## Discussion

*Nl*CCR, a robust Ca²⁺-permeable ChR, facilitates optical control of Ca²⁺-regulated cellular functions and is compatible with red-shifted fluorescent indicators. Compared with CapChR2, the best-performing of the previously available Ca^2+^-permeable ChRs^13^, *Nl*CCR generates larger and faster photocurrents, exhibits less desensitization and inward rectification, and displays higher Ca²⁺ selectivity; in addition, its blue-shifted action spectrum reduces crosstalk with red-shifted tools such as PinkyCaMP. By simultaneously monitoring PinkyCaMP response while activating *Nl*CCR, we confirmed that the ChR directly modulates intracellular Ca^2+^ levels in a light-dose-dependent manner. A previous analysis of the currents recorded from mouse hippocampal neurons upon CapChR2 photoactivation ^13^ revealed a substantial contribution of endogenous Ca^2+^-activated chloride channels (CaCCs). CaCCs are also expressed in HEK293 cells used in our study^28^, but their activation by increased cytoplasmic Ca^2+^ was unlikely in the presence of 10 mM EGTA in our intracellular solution. This conclusion is further corroborated by the rapid photocurrent kinetics in our experiments, which are consistent with direct *Nl*CCR activation.

Further development of optogenetic tools for control of intracellular Ca^2+^ requires an in-depth mechanistic understanding of *Nl*CCR function. As the first step towards this goal, we conducted a mutant screen and identified five residues in the putative cation-conducting pathway that are responsible for Ca^2+^ selectivity. Remarkably, three of these residues (Ser89, Asp96, and Glu233) are found in the positions critical for active proton transport by proton-pumping rhodopsins (Thr89, Asp96, and Asp212 in *Hs*BR)^29^. The residues corresponding to *Hs*BR’s Thr89 and Asp212 are conserved in most ChRs, but the presence of the Asp96 homolog is typical only of bacteriorhodopsin-like cation channelrhodopsins ^30^, to which *Nl*CCR shows only a low overall sequence homology. Gln238, highly conserved in ChRs, is adjacent to the Schiff base Lys237 and forms a part of the conserved “central gate”. Replacement of the corresponding residue with Glu increased the Ca^2+^ selectivity of *Co*ChR, from which CapChR2 was derived^13^. In contrast, the N238E mutation did not alter the Ca^2+^ selectivity of *Nl*CCR (Fig. 4a,b), suggesting that this conserved residue adopts different geometries and/or interacts with different partners in the two ChRs. Testing this hypothesis will require a high-resolution *Nl*CCR structure. Notably, except for the conservation of Gln238, there is little sequence homology between ancyromonad and chlorophyte ChRs in TM7 (Extended Data Fig. 1). Mutagenetic introduction of Glu residues at the other position in the central gate (the S21D mutaiton) increased the Ca^2+^ selectivity of *Nl*CCR (Fig. 4a,b), as the corresponding S43D mutation did in *Co*ChR^13^, suggesting a similar role of this residues in the two ChRs. The effect of the *Nl*CCR_M141C mutation in TM4 on selectivity can be explained by long-range interactions with pathway-forming residues in TM1-3 and TM7.

Overall, our mutational analysis identified several residues that determine *Nl*CCR Ca²⁺ permeability and show that this property can be further enhanced by introducing carboxylate residues into the central gate. However, the accompanying changes in photocurrent kinetics, rectification and amplitude indicate that Ca²⁺ permeability is coupled to multiple aspects of channel function. Thus, *Nl*CCR provides an engineerable scaffold for next-generation Ca²⁺-conducting ChRs, while highlighting the need to consider multiple ChR properties during optimization.

*Nl*CCR provides a genetically encoded means to directly manipulate intracellular Ca²⁺ with high spatiotemporal precision, enabling all-optical interrogation of Ca²⁺ dynamics^31^ and causal dissection of Ca²⁺-dependent signaling in intact neural circuits. By uncoupling Ca²⁺ influx from voltage-gated excitability, *Nl*CCR allows direct investigation of downstream pathways underlying neurotransmitter release, synaptic plasticity^32^, neurite outgrowth^33^, and other Ca²⁺-regulated neuronal processes across diverse cellular and network states. Because it does not rely on endogenous voltage-gated Ca²⁺ channels, this approach is also readily extendable to non-excitable cells, including glia, facilitating studies of Ca²⁺-dependent mechanisms implicated in neurological disease. More broadly, the central role of Ca²⁺ signaling throughout biology suggests applications extending well beyond neuroscience, including immune-cell activation^34^, excitation–contraction coupling in muscle, and many other Ca²⁺-regulated processes in development, physiology, and disease.

## Online content

Methods, additional references, Nature Research reporting summaries, statistical source data, supplementary tables, acknowledgments, details of author contributions and competing interests, a statement of data availability, and peer review information are available at…

## Acknowledgements

This work was supported by the National Institutes of Health grants R35GM140838 (J.L.S.), S10OD032293 (J.L.S.), U01NS118288 (M.X., J.L.S., F.S.P.), RF1NS133657 (J.L.S., F.S.P., M.X.), R61CA278458 (F.S.P.), and R01NS136027 (F.S.P.); the Robert A. Welch Foundation Endowed Chair AU-0009 (J.L.S.), and grants Q-2016-20220331 (F.S.P.) and Q-2016-20190330 (F.S.P.); a Vivian L. Smith Endowed Professorship in Neuroscience (F.S.P); the McNair Medical Foundation (F.S.P); M.X. is a Caroline DeLuca Scholar.

## Author contributions

EGG, OAS, FSP, MX, and JLS conceptualized the work and developed its methodology. EGG and OAS performed patch-clamp experiments in HEK293 cells, analyzed the results, and prepared the corresponding figures. HL and YW created *Nl*CCR and *Ans*ACR mutants and purified plasmid DNA for transfection. YG carried out neuronal experiments under the supervision of MX, and they both analyzed the results and prepared the corresponding figures. AJM designed, conducted, and analyzed the results of Ca^2+^ imaging experiments under FSP’s supervision; AJM and FSP prepared the corresponding figures. FSP, MX, and JLS provided the funds, supervised and administered the project. EGG, YG, AJM, and MX wrote an original draft, and all authors contributed to its review and editing.

## Competing interests

M.X. was a consultant to Capsida Biotherapeutics. Capsida Biotherapeutics provided research funds to Baylor College of Medicine to support a research project in the lab that is unrelated to this study and had no role in the research, authorship, and publication of this article.

## Methods

### Bioinformatics and molecular biology

The protein alignment was created using the MUSCLE algorithm with default parameters implemented in MegAlign Pro software v. 17.1.1 (DNASTAR Lasergene, Madison, WI) and truncated after the end of TM7. AlphaFold 3^35^ was used to create *Nl*CCR and CapChR2 homology models. PyMol (v. 3.1.8, Schrödinger) was used for molecular visualization.

For expression in human embryonic kidney (HEK293) cells, a mammalian codon-optimized polynucleotide encoding amino acid residues 1-272 of *Nl*CCR (Genbank Accession #PQ657779) was fused to a C-terminal mCherry tag and cloned into the pcDNA3.1(+) vector (Invitrogen, Cat. #V19520). The resultant plasmid is available from Addgene (#232600). The CapChR2 sequence encoding residues 1-288 was cloned from the Addgene plasmid #188032, fused to the same fluorescent tag, and cloned into the same vector backbone. The expression construct encoding residues 1-315 of *Cr*ChR2 (Genbank Accession #AF508966) was fused to a C-terminal EYFP (enhanced yellow fluorescent protein) tag and cloned into the same vector backbone. Point mutations were introduced using the QuikChange XL Site-Directed Mutagenesis Kit (Agilent, Cat. #200517) and verified by DNA sequencing. pCaggs-*Nl*CCR-TSER-mTurquoise2-ER-P2A-PinkyCaMP, pCaggs-CapChR2-TSER-mTurquoise2-ER-P2A-PinkyCaMP, and pCaggs-PinkyCaMP were assembled using the InFusion HD cloning kit (Takara Bio) and are available on Addgene (plasmid #’s). For neuronal expression, *Nl*CCR and CapChR2 genes were fused with EYFP and the Kv2.C-TlcnC membrane trafficking motif by PCR and NEBbuilder (NEB) to generate pAAV-CAG-*Nl*CCR-EYFP-Kv2.1C-TlcnC (Addgene #259919) and pAAV-CAG-CapChR2-EYFP-Kv2.1C-TlcnC (Addgene #259920), respectively.

### Automated whole-cell patch clamp recording from HEK293 cells

No cell lines from the list of known misidentified cell lines maintained by the International Cell Line Authentication Committee or non-human cell lines were used in this study. HEK293 cells were obtained from the American Type Culture Collection (ATCC, Cat. #CRL-1573), authenticated by short tandem repeats (STR) profiling at ATCC, and tested negative for mycoplasma contamination by PCR analysis. The cells were plated on 2-cm-diameter plastic dishes 48-72 hrs before the experiments, grown for 24 hrs, and transfected with 5 μl of Lipofectamine LTX with Plus Reagent (Thermo Fisher, Cat. #15338100) using 6 μg DNA per dish. All-*trans*-retinal (Millipore-Sigma, Cat. #116- 31-4) was added immediately after transfection at the final concentration of 5 µM.

Automated patch-clamp recordings were conducted at room temperature (21°C) using a SyncroPatch 384 (Nanion Technologies), as described earlier^15^. Transfected cells (48-72 h after transfection) were dissociated using TrypLE Express, diluted with CHO-S-SFM-II medium (both from ThermoFisher, Cat.# 12604013 and 31033020, respectively), and resuspended in External Physiological solution (Nation, Cat.# 08 3001). The chip was initially filled with a divalent-cation-free version of External Physiological solution to prevent blocking the holes with insoluble CaF_2_. After the addition of the cells and formation of GΩ-seals, this solution was replaced with External Physiological solution containing 2 mM CaCl_2_ and 1 mM MgCl_2_. We used NPC-384T 4-hole, S-type chips (Nanion, Cat. #222401) to increase the probability of capturing at least one cell expressing the transgene in each well. The photocurrent recorded from each well was the sum photocurrent of the four captured cells. To determine the V_r_ shifts upon replacement of Na^+^ with Ca^2+^, External Physiological solution was replaced with a solution containing 72 mM Ca^2+^ by serial dilution, and measurements were repeated in the same cells. The V_r_ values for other ionic conditions were determined by resuspending the cells in the corresponding solution and using the divalent-cation-free version of that solution to fill the chip. The holding voltages were corrected for LJPs. The full solution compositions and the corresponding LJP values calculated using the ClampEx LJP calculator are listed in Supplementary Table 1.

Illumination was provided with LUXEON Z Color Line light-emitting diodes (LEDs) Cat.# LXZ1-PB01 (470 ± 10 nm) arranged in a 6×16 matrix. The forward LED current was 900 mA (corresponding to an irradiance of ∼2 mW mm^-2^), the illumination duration was 200 ms (limited by the LED duty cycle), and the interval between successive light pulses was 60 s. The LEDs were driven by a derivative of CardioExcyte 96 SOL (Nanion, Cat. #191003) and controlled by Biomek commands. PatchControl384 v. 3.2 (Nanion Technologies) software was used to acquire data at a sampling rate of 5 kHz (200 μs per point). The current-voltage dependencies were recorded starting from -108.7 mV; the voltage was increased in 20-mV steps with 60-s dark intervals between successive recordings. The photocurrent amplitudes at the peak and the end of illumination, and the τ values derived from monoexponential approximation of photocurrent rise and decay were calculated using DataControl384 software v. 3.2 (Nanion Technologies). Further analysis was performed using OriginPro 2016 (OriginLab Corporation).

### Manual patch clamp recording from HEK293 cells

Manual patch clamp recordings to measure the photocurrent action spectra were performed with an Axopatch 200B amplifier (Molecular Devices). The pipette solution contained (in mM) KCl 130, MgCl_2_ 2, HEPES 10, pH 7.4, and the bath solution contained (in mM) NaCl 130, CaCl_2_ 2, MgCl_2_ 2, glucose 10, HEPES 10, pH 7.4. The signals were digitized with a Digidata 1440A (Molecular Devices) at a 5-kHz sampling rate (200 μs per point) using pClamp 10.7. Patch pipettes with 2-3 MΩ resistances were fabricated from borosilicate glass. Continuous light pulses were provided by a Polychrome V light source (T.I.L.L. Photonics GMBH) in combination with a mechanical shutter (Uniblitz Model LS6, Vincent Associates; half-opening time 0.5 ms). The action spectra of photocurrents were constructed by calculating the initial slope of photocurrent recorded in response to 15-ms light pulses at the irradiance <25 µW mm^-2^, corrected for the quantum density measured at each wavelength, and normalized to the maximal value.

### HEK293 cell culture and transfection in 96-well plates for Ca^2+^ imaging

To screen GEVIs for responses within the physiological range, we used HEK293 cells stably expressing the human Kir2.1 channel3, which maintains a resting membrane potential of approximately −83 mV using imaging solution #1 (in mM: NaCl 107, sucrose 26, glucose, HEPES 20, KCl 8.5, CaCl_2_ 2.5, MgSO_4_ 1.3, adjusted to pH 7.4 with NaOH, and adjusted to 300 mOsm/kg with H_2_O). This membrane potential was chosen because it approximates the lower bound of the physiological range for mammalian neurons, enabling screening for improved responses across the hyperpolarized state and into the depolarized range.

HEK293-Kir2.1 cells were cultured at 37 °C with 5% CO2 in growth medium #1 (high-glucose Dulbecco’s Modified Eagle Medium supplemented with 10% fetal bovine serum (FBS), 2 mM glutamine, 100 unit/mL penicillin, 100 μg/mL streptomycin, and 750 μg/mL of geneticin). Geneticin was included to maintain the expression of the Kir2.1 transgene, which was chromosomally integrated with a geneticin resistance gene. For screening GEVIs, glass-bottom 96-well plates (P96-1.5H-N, Cellvis) were first coated with poly-D-lysine (30-70 kDa) to promote cell adherence to the glass. The coating was performed for 1 hour at room temperature or 37 °C, and the plates were washed twice with DPBS. HEK293-Kir2.1 cells were then plated to 60-80% confluency in 100 μL of growth medium #2 (high-glucose Dulbecco’s Modified Eagle Medium supplemented with 5% FBS, 2 mM glutamine, 100 U/mL Penicillin, and 100 μg/mL Streptomycin).

Each of the three plasmids, pCaggs-*Nl*CCR-TSER-mTurquoise2-ER-P2A-PinkyCaMP, pCaggs-CapChR2-TSER-mTurquoise2-ER-P2A-PinkyCaMP, and pCaggs-PinkyCaMP, was transfected into 12 wells with jetPRIME (Polyplus) according to the manufacturer’s protocol. The following specifications were used: a mixture of 130 ng DNA, 0.4 μL jetPRIME transfection reagent, and 10 μL jetPRIME buffer was combined with 50 μL growth medium #2, then added to the well. Independent transfections were defined as those in which DNA was added to each well separately. After 6-18 hours, 100 μL growth medium #2 from each well was replaced with fresh growth medium #2 to minimize potential cytotoxicity from the transfection reagents. Two days post-transfection, the cells were washed twice with 200 μL of imaging solution at room temperature. Wells were filled with 100 μL of the imaging solution and assayed (see below). The imaging solution was adjusted with H_2_O to achieve a final osmolarity of 290-310 mOsm kg^-1^.

### Ca^2+^ fluorescence microscopy and image analysis

To evaluate changes in cellular Ca^2+^ concentrations induced by light-activated ChRs, we employed an all-optical assay on an inverted microscope (Ti, Nikon Instruments). To minimize spectral crosstalk, an illumination sequence in NIS-elements HC (version 5.42.06, Nikon Instruments) was constructed. Cells were exposed to 110 illumination cycles, each consisting of a 10 ms pulse of 7.4 mW mm^-2^ 575/25 nm light (central wavelength/bandwidth) from a SpectraIII (Lumencor), followed by 490 ms of darkness. In the 11^th^ cycle, in addition to the 575/25 nm pulse, a concurrent 440/20 nm pulse was added to activate the ChR. Time-lapses (∼55 seconds) from four separate fields of view (FOVs) were collected for each well with the same 575/25 nm pulse pattern and varying 440/20 nm pulse irradiance (0, 0.9, 4.4, and 17.6 mW mm^-2^). Time-lapses were collected using a Kinetix sCMOS camera (Teledyne Vision Solutions) mounted on a 0.45× demagnifying tube (Nikon). Each 10-ms exposure frame of the time-lapse consisted of a 16-bit, 960 x 960-pixel image collected through a 5-band (432/515/595/681/809 nm) filter cube (part# 77015970, Semrock).

For single-cell analysis, frames corresponding to open imaging illumination paths were first compiled into time-series image stacks. Frames acquired during activation illumination were excluded from downstream analysis due to spectral bleed-through of the activation light through the optical filter cube. This produced a substantial artifactual increase in the detected signal, independent of cellular fluorescence, and therefore could not be reliably used for quantitative analysis. Time-series image stacks were then max-intensity projected to generate an image for cellular segmentation using Cellpose3^36^. Segmentation masks generated for each FOV were then applied to the full time-series dataset to extract fluorescence traces per cell. For each segmented cell, the baseline fluorescence intensity *F*_0_ was calculated from the pre-stimulation period, and fluorescence responses were recorded as Δ*F*/*F*_0_across the entire recording. To control for optical crosstalk and non-specific activation arising from the imaging illumination, recordings acquired under no-activation-pulse conditions were used as baseline control traces. These control traces were subtracted to isolate fluorescence changes specifically associated with the intended stimulation pulse and subsequent Ca^2+^ channel activation.

To account for potential variability in Ca^2+^ channel expression levels across cells, fluorescence responses were additionally normalized to the intensity of the coexpressed reference channel, yielding an expression-normalized response metric. For each quantified metric, responses from all segmented cells within a given well were averaged to generate a single well-level response value for statistical analysis. All image processing, segmentation, signal extraction, and quantitative analyses were implemented in Python 3.12.10 using custom written analysis software. The excitation spectrum of PinkyCaMP was adapted from ^16^ with the authors’ permission.

### Mice

All procedures to maintain and use mice were approved by the Institutional Animal Care and Use Committee at Baylor College of Medicine (protocol AN-6544). Mice were maintained on a 14 hr:10 hr light:dark cycle with regular mouse chow and water *ad libitum*. The temperature was maintained at 21–25°C and a humidity at 40-60%. Experiments were performed during the light phase. Female ICR (CD-1) mice were purchased from Baylor College of Medicine Center for Comparative Medicine, and male C57BL6/J (JAX #000664) mice were obtained from Jackson Laboratory. Both male and female mice were used in the experiments.

### *In utero* electroporation

Female ICR mice were crossed with male C57BL6/J mice to obtain timed pregnancies. *In utero* electroporation was used to deliver the transgenes ^37^. To express *Nl*CCR-EYFP-Kv2.1C-linker-TlcnC (*Nl*CCR-K-T), CapChR2-EYFP-Kv2.1C-linker-TlcnC (CapChR2-K-T), or *Cr*ChR2_H134R-EYFP in the layer 2/3 pyramidal neurons of the somatosensory cortex, plasmids pAAV-CAG-*Nl*CCR-EYFP-Kv2.1C-linker-TlcnC, pAAV-CAG-CapChR2-EYFP-Kv2.1C-linker-TlcnC, or pCAG-hChR2_H134R-EYFP (Addgene #114367) (2.5 μg μl^-1^ final concentration) was mixed with pCAG-tdTomato (0.1 μg μl^-1^ final concentration) and Fast Green (Sigma-Aldrich, 0.01% final concentration) for injection. On embryonic day 15, pregnant mice were anesthetized, and a beveled glass micropipette (tip size 100-μm outer diameter, 50-μm inner diameter) was used to penetrate the uterus and the embryonic skull to inject ∼1.5 μl DNA solution into one lateral ventricle. Five pulses of current (voltage 39 V, duration 50 ms) were delivered at 1 Hz with a Tweezertrode (5-mm diameter) and a square-wave pulse generator (Gemini X2, BTX Harvard Bioscience). The electrode paddles were positioned in parallel with the brain’s sagittal plane. The cathode was placed in contact with the side of the brain ipsilateral to the injected ventricle to target the somatosensory cortex. Transfected pups were identified by tdTomato fluorescence via transcranial imaging with an MZ10F stereomicroscope (Leica) 1 day after birth.

### Brain slice electrophysiology and imaging

Mice were used at 3-6 weeks of age for acute brain slice electrophysiology experiments. Mice were anesthetized by an intraperitoneal injection of a ketamine and xylazine mix (80 mg kg^-1^ and 16 mg kg^-1^, respectively) and perfused transcardially with cold (0–4°C) slice cutting solution containing 80 mM NaCl, 2.5 mM KCl, 1.3 mM NaH_2_PO_4_, 26 mM NaHCO_3_, 4 mM MgCl_2_, 0.5 mM CaCl_2_, 20 mM *d*-glucose, 75 mM sucrose and 0.5 mM sodium ascorbate (315 mOsm l^-1^, pH 7.4, saturated with 95% O_2_/5% CO_2_). Brains were dissected and sectioned in the cutting solution using a VT1200S vibratome (Leica) to obtain 300-μm coronal slices. Slices were incubated in a custom-made interface-holding chamber containing slice-cutting solution saturated with 95% O_2_/5% CO_2_ at 34°C for 30 min, and then at room temperature for 20 min to 8 h until they were transferred to the recording chamber. We performed recordings on submerged slices in artificial cerebrospinal fluid (ACSF) containing 119 mM NaCl, 2.5 mM KCl, 1.3 mM NaH_2_PO_4_, 26 mM NaHCO_3_, 1.3 mM MgCl_2_, 2.5 mM CaCl_2_, 20 mM *d*-glucose and 0.5 mM sodium ascorbate (305 mOsm l^-1^, pH 7.4, saturated with 95% O_2_/5% CO_2_, perfused at 3 ml min^-1^) at 30–32°C. Neurons were visualized with video-assisted IR differential interference contrast imaging, and fluorescent neurons were identified by epifluorescence imaging under a water immersion objective (×40, 0.8 NA) on an upright SliceScope Pro 1000 microscope (Scientifica) with an IR-1000 CCD camera (DAGE-MTI). For whole-cell recordings, a K^+^-based pipette solution containing 142 mM K^+^ gluconate, 10 mM HEPES, 1 mM EGTA, 2.5 mM MgCl_2_, 4 mM ATP-Mg, 0.3 mM GTP-Na, 10 mM Na_2_-phosphocreatine (295 mOsm l^-1^, pH 7.35) or a Cs^+^-based pipette solution containing 121 mM Cs^+^-methanesulfonate, 10 mM HEPES, 10 mM EGTA, 1.5 mM MgCl_2_, 4 mM ATP-Mg, 0.3 mM GTP-Na, 10 mM Na_2_-phosphocreatine, and 2 mM QX314-Cl (295 mosmol, pH 7.35) was used to record photocurrent of ChR^+^ neurons or EPSCs of ChR^−^ neurons, respectively. Data were acquired at 10 kHz and low-pass filtered at 4 kHz with an Axon Multiclamp 700B amplifier and an Axon Digidata 1440A Data Acquisition System under the control of Clampex 10.7 (Molecular Devices). Data were analyzed offline using ClampFit 10.7 (Molecular Devices) and AxoGraph 1.7.6. For photostimulation, blue light was emitted from a collimated 470-nm light-emitting diode (LED; M470L3, Thorlabs) to activate ChRs. The LEDs were driven by an LED driver (Throlabs LEDD1B) under the control of an Axon Digidata 1440A Data Acquisition System and Clampex 10.7. The light was delivered through the reflected light fluorescence illuminator port and the 40× objective.

To measure the photocurrent of ChRs, ChR^+^ neurons were clamped at -70 mV with the K^+^-based internal solution, and 100-ms 470-nm (38.7 mW mm^-2^) light stimulation was applied. To test whether photocurrents were altered by antagonists, the same light was used to activate CapChR2 and *Cr*ChR2_H134R while light power was reduced to 1.8 µW mm^-2^ to stimulate *Nl*CCR to reduce photocurrent and error caused by series resistance. To record EPSCs evoked by activating ChRs, ChR^-^ neurons were clamped at -60 mV with the Cs^+^-based internal solution and 100-, 10-, or 1-ms 470-nm (38.7 mW mm^-2^) light stimulation was applied. The drugs were added to the ACSF at the following concentrations: TTX (1 μM), TEA (1.5 mM), 4-AP (1.5 mM), CdCl_2_ (0.1 mM), NBQX (10 μM), and (RS)-CPP (10 μM).

After electrophysiology recordings, fluorescent images of the brain slices were acquired on an Axio Zoom V16 Fluorescence Stereo Zoom Microscope (Zeiss) and processed using MATLAB2024b (MathWorks). Images were taken from 21 brain slices of one male and three female mice.

### Statistics and reproducibility

Plasmids encoding different ChR variants were randomly assigned to identical batches of HEK293 cells for transfection. In automated patch-clamp studies, cells were blindly selected by the machine and randomly distributed to the wells. Data from the 4-hole wells that formed seals with resistance <100 MΩ were excluded from the analysis. Also disregarded were wells yielding end photocurrents with absolute magnitude <100 pA at -100 mV, to exclude cells expressing no transgene and improve the signal-to-noise ratio for more accurate estimation of the V_r_ values by plotting the current-voltage dependencies. Traces on which the curve fitting algorithm failed to converge were excluded from the estimation of the kinetic parameters. Measurements were replicated across four independent batches of cells transfected with wild-type *Nl*CCR to compare it with other ChR variants, and across three batches to compare each *Nl*CCR mutant with the wild type. Data from all batches transfected with the same variant were pooled. Unless otherwise stated (e.g., in experiments with external solution exchange), only one photocurrent trace was recorded per cell and condition. Photocurrent traces recorded from different wells transfected with the same transgene were treated as biological replicates (reported as n values). These values indicate how often the experiments were performed independently.

Statistical analysis of HEK293 patch-clamp data was performed using OriginPro 2016. To compare data obtained under different ionic conditions, the normal distribution assumption was not made, and the non-parametric two-tailed Mann-Whitney test was used to compare the means. The normality of the data obtained with the standard external solution was assessed using the Kolmogorov-Smirnov test, and the data were analyzed using one-way ANOVA followed by Tukey’s test for pairwise comparisons of the means. The number of wells sampled (n values) is indicated in the figures or figure captions or provided, along with the numerical raw data, in the Source Data Files. No statistical methods were used to pre-determine sample sizes, but the sample sizes were within the ranges reported in previous publications^7, 38^.

For Ca^2+^ imaging experiments, n = 12 wells were independently transfected for each construct. One well of *Nl*CCR pulsed with 17.6 mW mm^-2^ light was removed due to a hardware failure. Cells (>100) in each FOV were individually analyzed for their Ca^2+^ responses, and the resulting traces were averaged. Statistical analyses were performed on pooled FOV-level data by Prism 11 (Graphpad). A Shapiro-Wilk test showed that the data in Fig. 2g were lognormal, with P values of 0.86, 0.66, and 0.59 for *Nl*CCR, CapChR2, and no-ChR2 control, respectively. A lognormal ordinary one-way ANOVA revealed a significant effect of group on peak response (F(3, 44) = 2160, P = 9.2 × 10^⁻48^, η² = 0.99). Dunnett’s multiple comparisons test revealed that the peak response of NlCCR was significantly greater than that of CapChR2 (P = 4.1 × 10^⁻14^) and the no-ChR control (P < 1 × 10^⁻15^). All statistical tests were two-sided.

Statistical analysis of the brain slice data was performed by Prism 11 (Graphpad). Data were tested for normality. For data passing the normality tests, t test for 2 groups and one-way ANOWA for more than 2 groups were applied. For data not passing the tests, nonparametric tests were applied. Both male and female were included in the study. The number of animals sampled is indicated in the figure captions along with the numerical raw data in the Source Data Files. No statistical methods were used to predetermine sample sizes.

## Data availability

The numerical data and statistical analyses are provided in Source Data Files. The plasmids *Nl*ChR_mCherry_pcDNA3.1, pCaggs-*Nl*CCR-TSER-mTurquoise2-ER-P2A-PinkyCaMP, pCaggs-CapChR2-TSER-mTurquoise2-ER-P2A-PinkyCaMP, pCaggs-PinkyCaMP, pAAV-CAG- *Nl*CCR-EYFP-Kv2.1C-linker-TlcnC, and pAAV-CAG-CapChR2-EYFP-Kv2.1C-linker-TlcnC are available from Addgene. All other data and materials are available to any researcher upon reasonable request, subject to materials transfer agreements (MTAs).

## Extended Data figures and captions

**Extended Data Figure 1.**
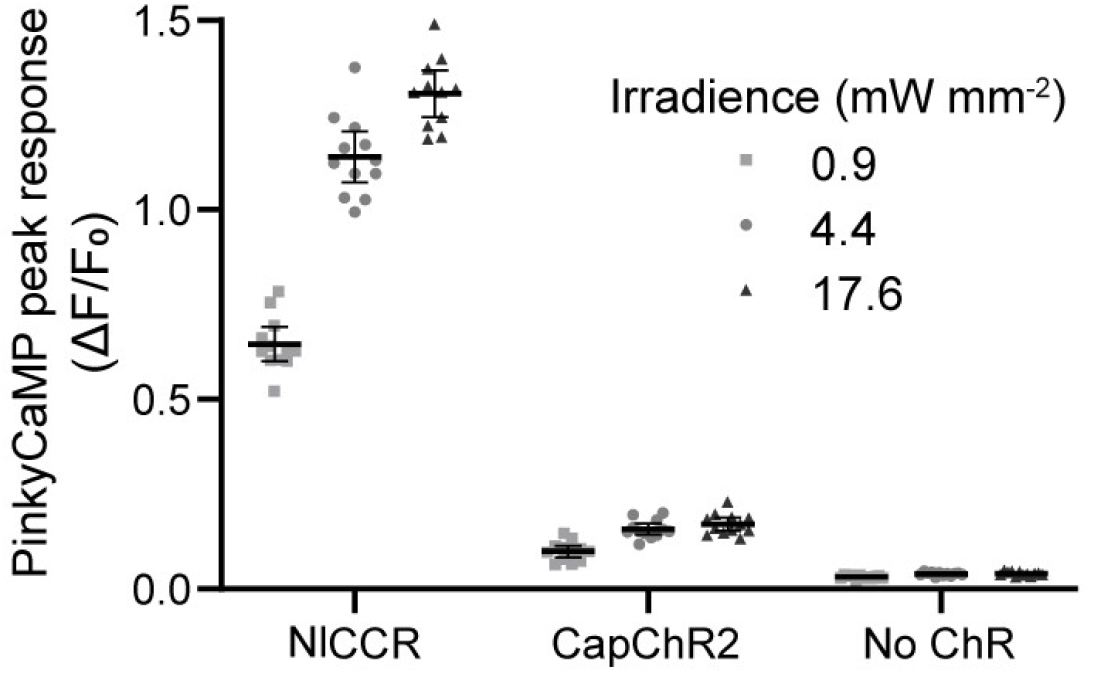
Individual data points for the peak responses shown in Fig. 2f. PinkyCaMP response to ChR (or no ChR control) activation by 440/20 nm light expressed in HEK293 Kir2.1 cells. n = 12 independently transfected wells.

**Extended Data Figure 2.**
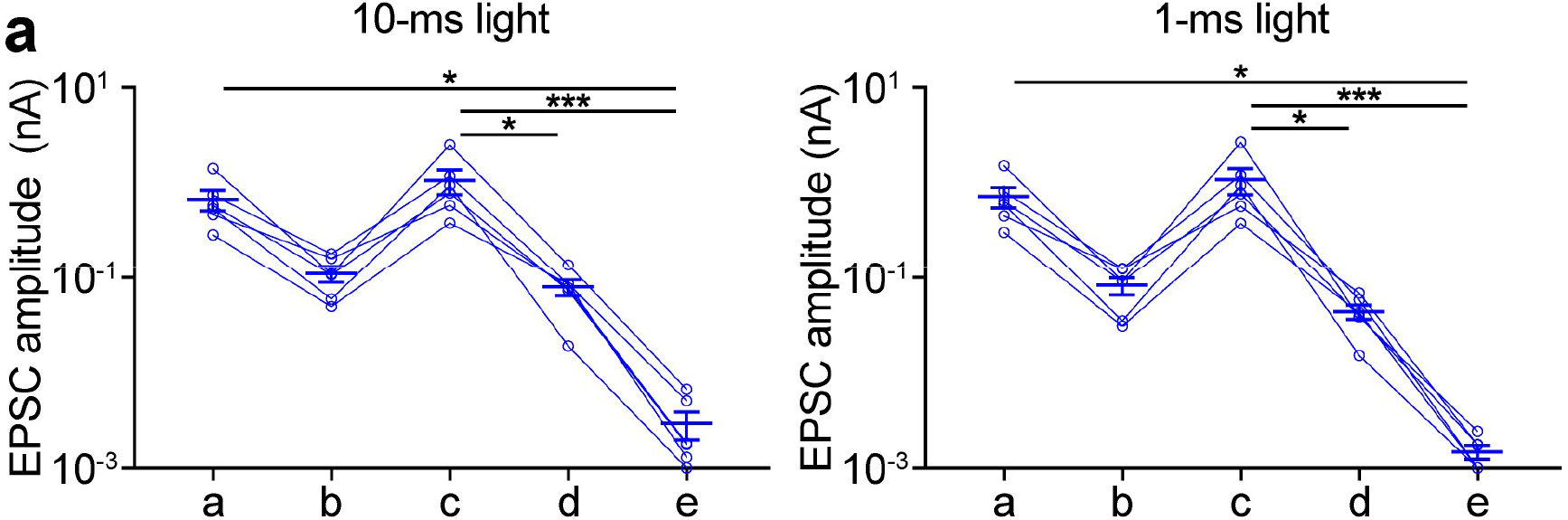
*Nl*CCR enables neurotransmitter release without the activation of VGCCs. (**a**) Summary data of evoked EPSC peak amplitudes of recorded *Nl*CCR^−^ neurons (*n* = 6) in the brain slices expressing *Nl*CCR in response to 10-ms (left) or 1-ms (right) 470-nm light stimulation under the indicated conditions as in Figure 3d. In all panels, each symbol represents a recorded neuron. Data are from 2 female mice. Summary data are expressed as mean ± SEM. *, P < 0.05; ***, P < 0.001 by Friedman test with Dunn’s correction.

**Extended Data Figure 3.**
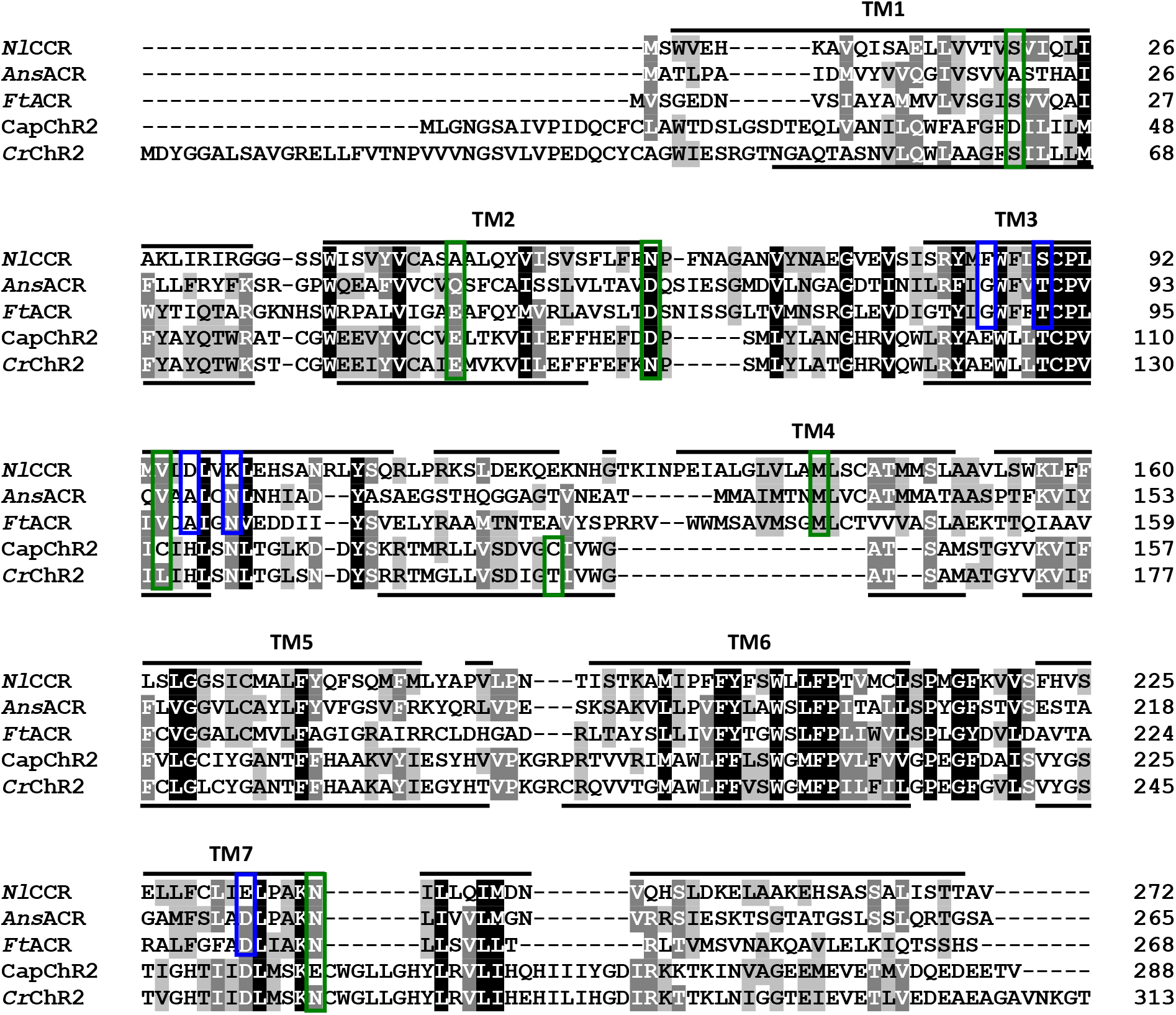
Protein alignment of the transmembrane (7TM) domains, shaded according to the degree of residue conservation. The black lines at the top of the alignment indicate the α-helical regions of the *Nl*CCR homology model; those at the bottom indicate the α-helical regions of the *Cr*ChR2 structure (6EID). The residue positions in which the *Nl*CCR sequence diverges from the *Ans*ACR and *Ft*ACR sequences, tested by the mutations, are boxed blue; those in which the *Nl*CCR residues were mutagenetically replaced with the CapChR2 residues are boxed green. The numbers on the right are the last residue numbers in each line.

**Extended Data Figure 4.**
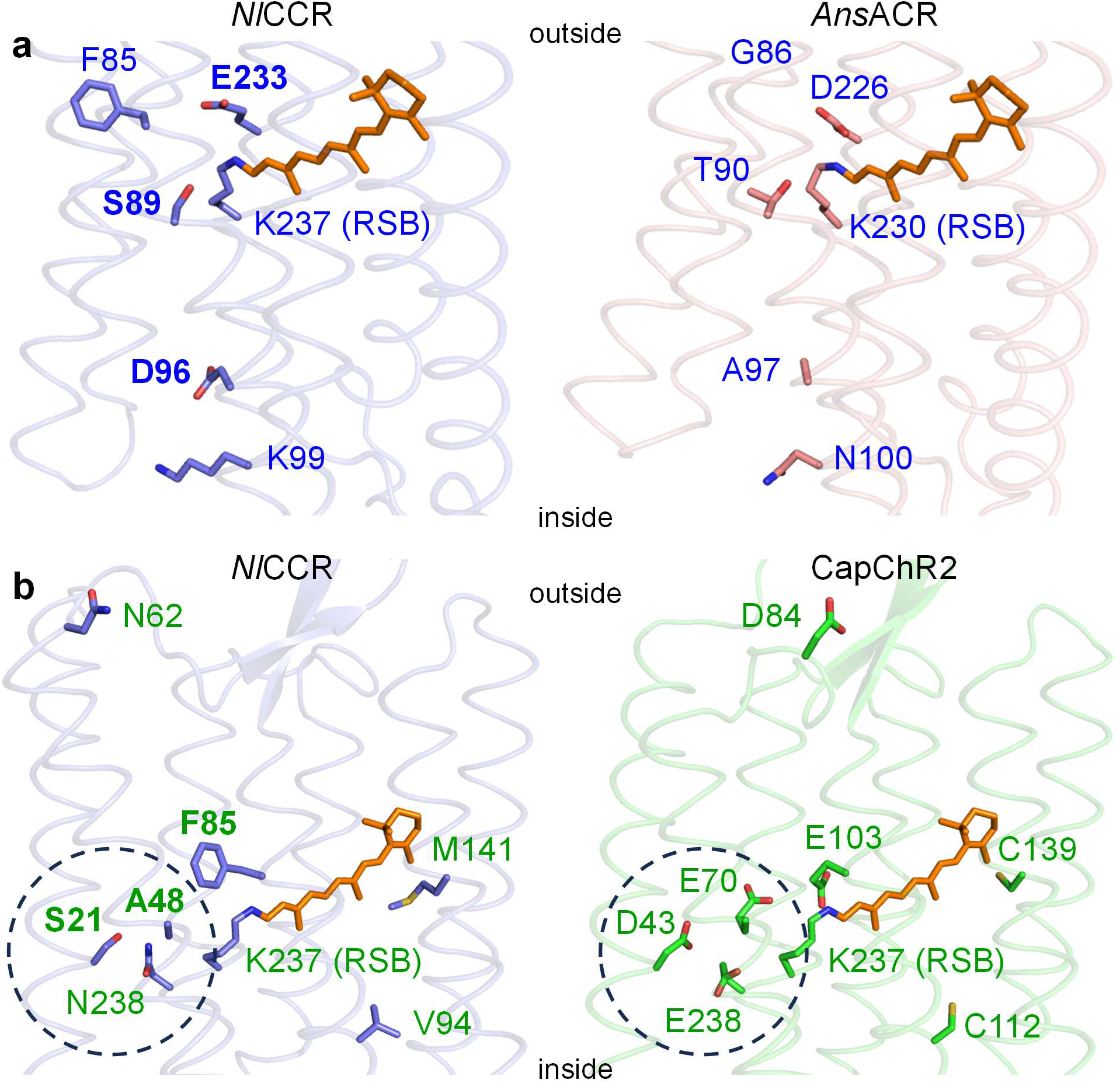
**a**, Homology models of *Nl*CCR (blue) and *Ans*ACR (pink) showing the side chains of the tested divergent residues. **b**, Homology models of *Nl*CCR (blue) and CapChR2 (green) showing the side chains of the tested divergent residues. The retinal chromophore is shown in orange. The dashed circle indicates the residues that form the central gate. RSB, retinal Schiff base. Bold labels mark the side chains that determine Ca^2+^ selectivity of *Nl*CCR.

**Extended Data Figure 5.**
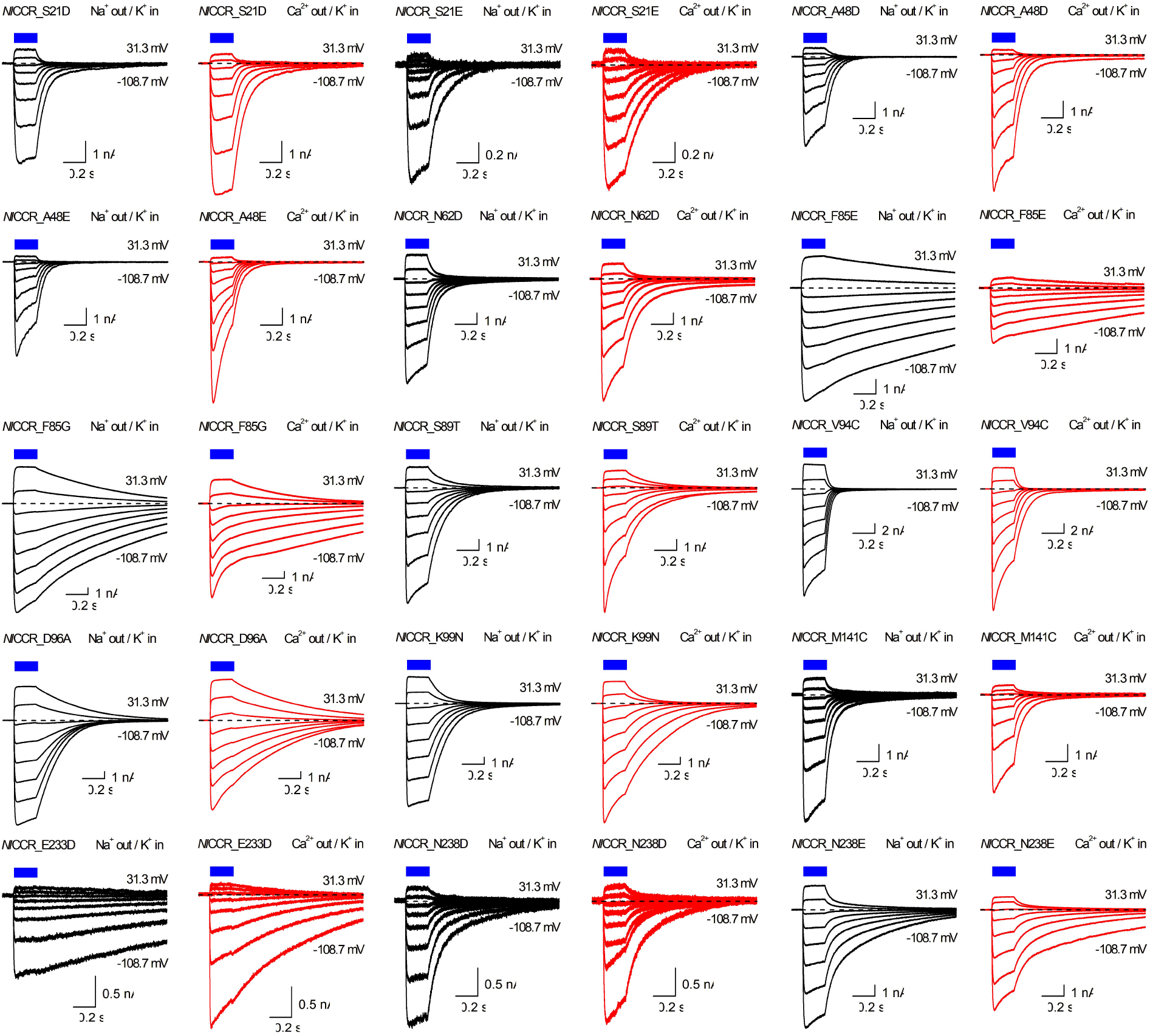
Photocurrent traces recorded from the indicated *Nl*CCR mutants upon incremental voltages in the Na^+^-based (black) or Ca^2+^-based (red) external solution in response to 200-ms light pulses, the duration of which is shown by the blue rectangles. The holding voltages shown in the plots are corrected for LJPs.

**Extended Data Figure 6.**
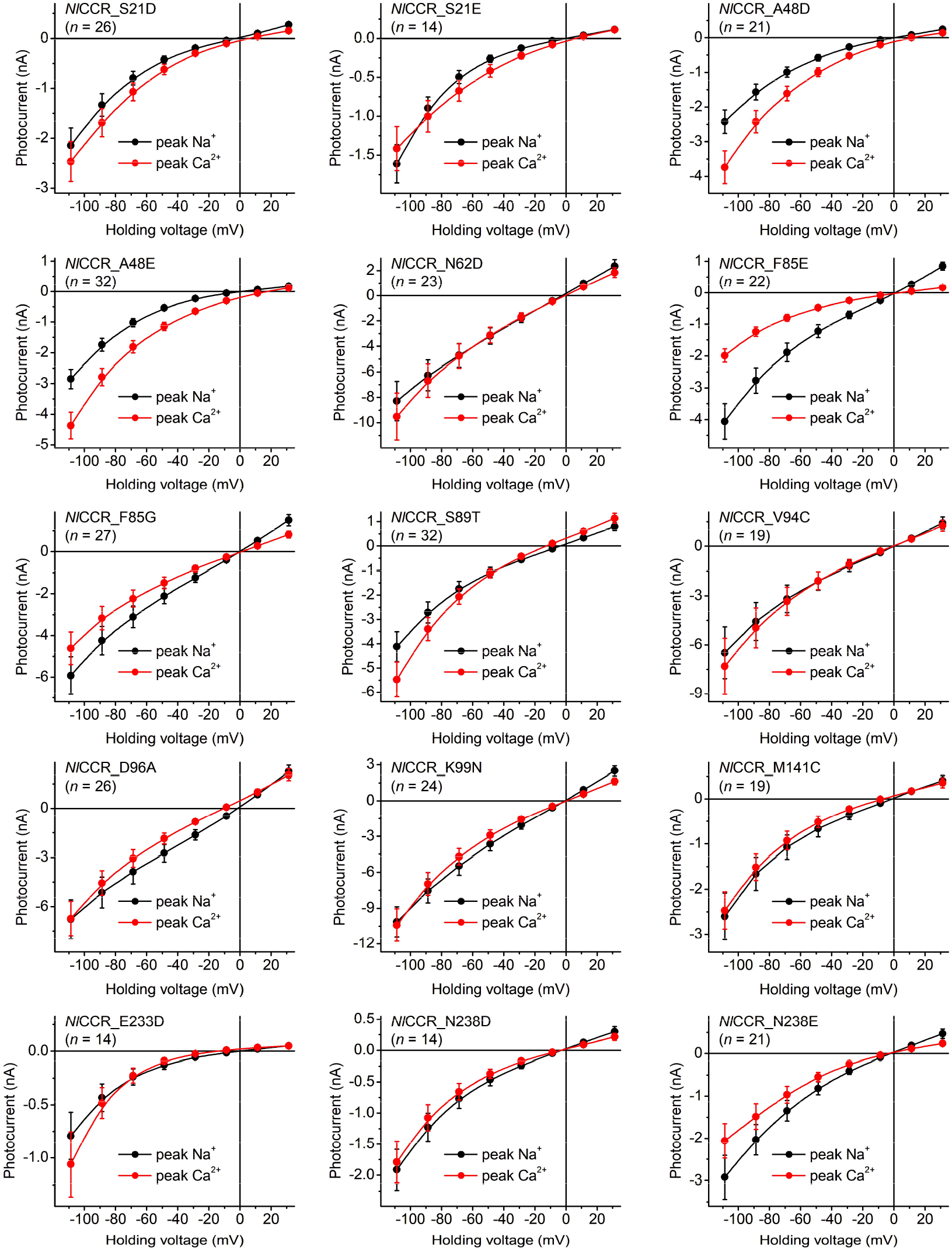
The voltage dependencies of the peak photocurrents recorded from the *Nl*CCR mutants in the Na^+^-based external solution (black) and after its exchange to the Ca^2+^-based external solution (red). The symbols represent mean ± SEM values; the numbers of wells sampled per variant (n) are shown in the plots.

**Extended Data Figure 7.**
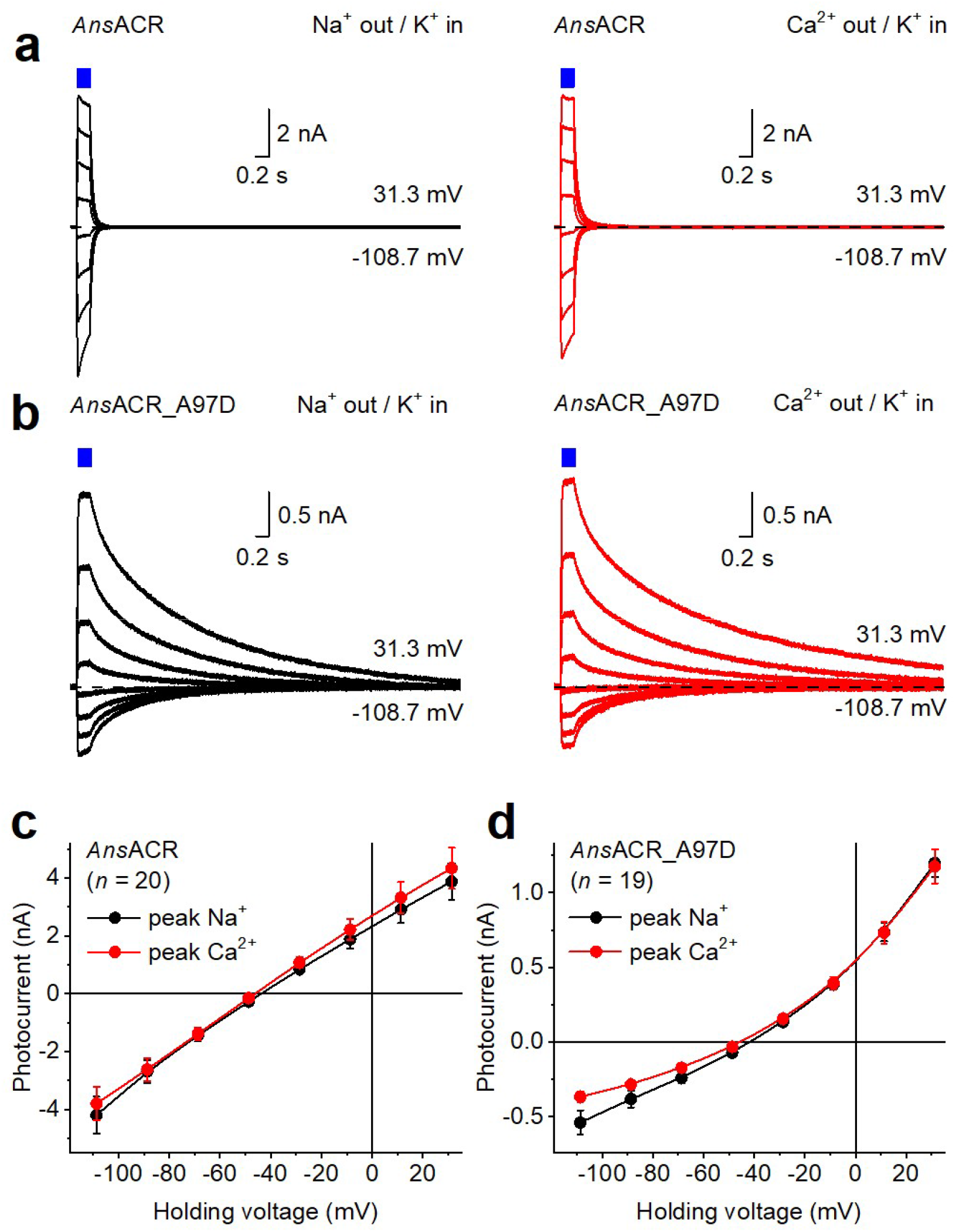
**a,b**, Photocurrent traces recorded from the wild-type *Ans*ACR and its A97D mutant upon incremental voltages in the Na^+^-based (black) or Ca^2+^-based (red) external solution in response to 200-ms light pulses, the duration of which is shown by the blue rectangles. The holding voltages shown in the plots are corrected for LJPs. **c,d**, The voltage dependencies of the peak photocurrents recorded from the wild-type *Ans*ACR and its A97D mutant in the Na^+^-based external solution (black) and after its exchange to the Ca^2+^-based external solution (red). The symbols represent mean ± SEM values; the numbers of wells sampled per variant (n) are shown in the plots.

