## Supplementary Table 1 for "A highly Ca^2+^-permeable channelrhodopsin from the ancyromonad *Nutomonas longa* enables optogenetic control of Ca^2+^ signaling"

**Supplementary Table 1. Compositions of the solutions used in automated patch-clamp recording in HEK293 cells**

| | KF | NaCl | KCl | NMDG Cl | $CaCl_2$ | $MgCl_2$ | HEPES | EGTA | glucose | LJP |
|---|---|---|---|---|---|---|---|---|---|---|
| **Internal solution pH 7.2** | 110 | 10 | 10 | — | 2 | 1 | 10 | 10 | — | — |
| **Standard external solution pH 7.4** | — | 140 | 4 | — | 2 | 1 | 10 | — | 5 | 8.7 |
| **$CaCl_2$ external solution** | — | — | 4 | — | 72 | 1 | 10 | — | 5 | — |
